# *RADIALIS* origin and expression suggest ancestral function in female organs of seed plants

**DOI:** 10.1101/2025.07.30.667799

**Authors:** Aniket Sengupta, Dianella G. Howarth

## Abstract

**Premise of the study:** Pairs of protein homologs may participate in competitive interactions to define morphology. How these competitive pairs evolve, whether they evolve repeatedly, and how they affect the origin of novel features are open questions. Ovules/seeds are a major innovation in plants, a trait that evolved in the ancestor of all seed plants. Within seed plants, flowers and fruits are synapomorphies arising within angiosperms, while gymnosperms retain the ancestral absence of these structures. RADIALIS and DIVARICATA are two MYB homologs whose competitive interaction is involved in fruit/carpel development and flower symmetry.

RADIALIS have a protein-binding domain, but DIVARICATA have an additional DNA-binding domain.

**Methods:** We reconstructed the phylogenetic history of these genes with a Bayesian approach. We tested the expression of these genes in *Ginkgo biloba* with real-time PCR.

**Key results:** We demonstrate that *DIVARICATA* genes underwent two rounds of duplications at the base of vascular plants forming three clades: *DIV-A*, *DIV-B*, and *DIV-C*. We show that *RADIALIS* homologs evolved only once: from *DIV-C* at the base of seed plants, mediated by a premature stop codon likely generated by a single-base substitution. We surveyed the expression pattern of these genes for the first time in a gymnosperm, *Ginkgo biloba*. We find that *Ginkgo biloba RADIALIS* genes often have higher expression in ovules. This is consistent with the expression and function of RADIALIS in angiosperm carpels.

**Conclusions:** Our work provides suggestive evidence that the evolution of seed habit may be associated with the origin of the silencing peptide RADIALIS.

**Significance Statement:** We determine the origin of *RAD* genes from *DIV* genes near the base of seed plants through truncation caused by an early stop codon. We found that the *RAD* genes of the early-diverging gymnosperm *Ginkgo biloba* have a high expression in the female organs, similar to that of many flowering plants, suggesting a possible role of RAD genes in female organs across seed plants.

## Introduction

An ongoing question in biology is how gene duplication and changes in gene expression may be associated with morphological shifts. Gymnosperms and angiosperms make up the clade spermatophyta, or seed plants, and are characterized by the evolution of seeds. In order to understand how certain genes were recruited to petal and ovary development in angiosperms, we need to analyze these genes in gymnosperms that preserve the ancestral state of the absence of flowers. Thus, seed plants provide a good system to understand how gene duplication and changes in gene expression may be associated with evolution of novel morphology, like that of flowers.

Flowers are complex structures whose development is under the control of several gene families—including the *MYB* family (reviewed in the following: Ma *et al*., 2017; Wu *et al*., 2022; Wang *et al*., 2023). The function of MYB proteins is well-understood in flower symmetry (Galego & Almeida, 2002; Corley *et al*., 2005; Raimundo *et al*., 2013; Su *et al*., 2017), which has been studied mostly in context of the corolla whorl. Ancestral flowers were radially symmetrical, with bilaterally symmetric flowers evolving at least 130 times independantly during the diversification of flowering plants (Reyes *et al*., 2016). Bilaterally symmetric flowers often have a distinct morphology from the dorsal to the ventral side. The control of bilateral symmetry in flowers by MYB proteins was first reported in *Antirrhinum majus* (snapdragons, order Lamiales) where the MYB protein *A. majus* RADIALIS (AmRAD) defines the dorsal side of the flower (Corley *et al*., 2005) and the MYB protein *A. majus* DIVARICATA (AmDIV) defines the ventral side (Galego & Almeida, 2002). Furthermore, there is growing evidence that RAD homologs are also involved in the development of the complex female structures inside flowers called gynoecia or carpels, across flowering plants (Sengupta & Hileman, 2022; Masuda *et al*., 2022; Cheng *et al*., 2023). In contrast, gymnosperms do not have flowers, therefore do not have petals or gynoecia. However, gymnosperms have *RAD* and *DIV* genes (Raimundo *et al*., 2018), and also have simpler reproductive organs. This system is suited for comparative analysis on how the duplication history and the expression pattern of *RAD* and *DIV* in gymnosperms can inform the origin of flowers in angiosperms. *RAD* genes have only been found in angiosperms and gymnosperms, but *DIV* genes have been reported from all groups of green plants (Raimundo *et al*., 2018).

AmDIV proteins have two MYB domains (Lucibelli *et al*., 2020)—MYB1 and MYB2. The MYB1 domain is present near the N-terminus (5′-end of the mRNA), and is predicted to be a protein-binding SANT domain, whereas and the MYB2 domain located near the C-terminus domain is predicted to be either a SANT (Lu *et al*., 2020) or a HTH myb-type (Sigrist *et al*., 2013; ‘DIVARICATA in UniProt’). AmDIV functions as a transcription factor (Raimundo *et al*., 2013). It is likely that this DNA-binding activity of AmDIV is performed by the MYB2 domain; this argument is based on indirect studies from homologs—both artificial (Rose *et al*., 1999) and natural (Raimundo *et al*., 2013)—lacking the MYB2 domain. Further, to be able to function, AmDIV needs to heteromerize with two co-transcription factors called *A. majus* DIVARICATA AND RADIALIS INTERACTING FACTOR 1 and 2 (AmDRIF1 and AmDRIF2) (Raimundo *et al*., 2013) by likely employing its MYB1 domain. The AmRAD protein, on the other hand, has been reported to have a MYB1 domain (but not the MYB2 domain), and can also bind to DRIF1/2 proteins (Raimundo *et al*., 2013).

AmRAD therefore acts as a competitive inhibitor of the AmDIV protein: it can bind to AmDRIF1/2 (because it has a MYB1 domain) but cannot act as a transcription factor (because it has no MYB2 domain) (Corley *et al*., 2005; Barg *et al*., 2005; Machemer *et al*., 2011). Such regulatory proteins have been described as short interfering peptides (siPEPs) (Seo *et al*., 2011; Bartlett & Whipple, 2013), microProteins (miPs) (Staudt & Wenkel, 2011; Magnani *et al*., 2014), and small regulatory proteins (SRPs) (Floyd *et al*., 2014). *RAD* has been described as a truncated version of *DIV* (Davies *et al*., 2006; Howarth & Donoghue, 2009), and has been hypothesised to have originated from *DIV* either by alternative splicing or by partial deletion (Raimundo *et al*., 2018). Truncated miPs that evolved through gene duplication followed by truncation of the coding sequence are called trans-miPs (Magnani *et al*., 2014). If *RAD* indeed is a truncated version of *DIV*, some process that may explain this include segmental deletion, evolution of a premature stop codon by substitution, insertion of a transposon, or changes in intron-exon boundaries.

RAD-DIV competitive interaction has been described from two groups of asterid angiosperms: Lamiales and Solanales (Machemer *et al*., 2011; Raimundo *et al*., 2013). This interaction has been suggested to be homologous between these two orders (Sengupta & Hileman, 2022). However, *RAD*, *DIV*, and *DRIF* genes are present across a much larger group of plants than Lamiales and Solanales, and display interactions in yeast-two-hybrid assays (Raimundo *et al*., 2018), suggesting that a RAD-DIV competitive interaction is possibly widespread. Yet, the phylogenetic relationship between *DIV* and *RAD*, and the origin of the RAD-DIV interaction remain unresolved. For example, it is not clear whether there have been parallel origins of *RAD* or *DIV* by loss/gain of the *MYB2* domain in various species (Fig. 1). At least one case of independent evolution of truncated siPEP/miP has been reported: truncated siPEP/miP targeting the transcription factors Meis/KNOX evolved independently in flowering plants and *Caenorhabditis elegans* (Magnani *et al*., 2014). On the other hand, the truncated LITTLE ZIPPER proteins that target class III Homeodomain Leucine Zipper (C3HDZ) proteins evolved once at the base of ferns + seed plants via loss of DNA-binding domain (Floyd *et al*., 2014).

**Fig. 1.**
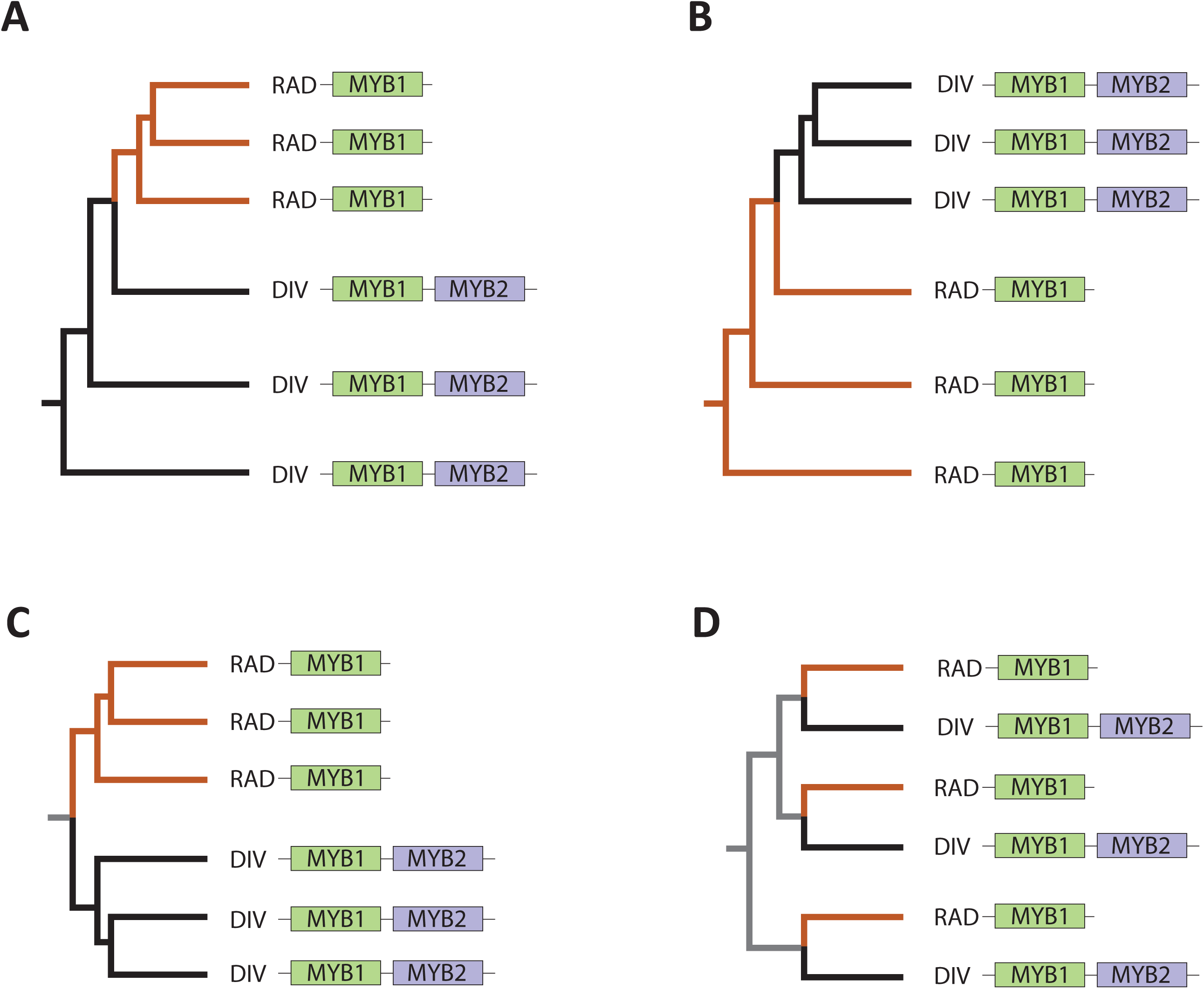
Some possible hypotheses that may explain the origin of RAD and DIV proteins and their interaction. More complex hypotheses are possible but not represented here. **A.** *RAD* evolved from *DIV* through truncation if MYB2 domain, hence, is nested within *DIV*. The common ancestor had domain organization similar to a *DIV* gene. **B**. *DIV* evolved from *RAD* through gain of MYB2 domain, hence is nested within *RAD*. The common ancestor had domain organization similar to a *RAD* gene. **C.** *RAD* and *DIV* are reciprocally monophyletic. The common ancestor had domain organization that was different from either *DIV* or *RAD*. **D.** Either *RAD*-like genes evolved multiple times from *DIV* genes, or *DIV*-like genes evolved multiple times from *RAD* genes. Black branches represent a *DIV*-like ancestral state, orange branches represent a *RAD*-like ancestral state, grey branches represent ambiguity.

These hypotheses regarding the origin of the RAD-DIV interaction can be tested by identifying *RAD* and *DIV* genes from a wide variety of green plants and reconstructing a phylogenetic tree. Indeed, not just for RAD and DIV, these hypotheses remain untested for many pairs of siPEP and their target genes (Bartlett & Whipple, 2013). We test the relationship among *DIV* and *RAD* genes from various species by employing a Bayesian Phylogenetic approach. We also test the expression of these genes in *Ginkgo biloba* using quantitative real-time PCR (qRT-PCR). *Ginkgo biloba* is a monotypic gymnosperm lineage, that is sister to cycads; together, they are sister to the rest of the extant gymnosperms (Liu *et al*., 2022; Yang *et al*., 2022). *Ginkgo* are dioecious and the female trees produce seeds that develop fleshy sacrotesta enclosing a hard sclerotesta (giving the seeds a drupe-like appearance). This makes *G. biloba* a good system to understand the broader gene expression patterns of *RAD* and *DIV* genes across seed plants. We find that *RAD* genes evolved from *DIV* genes at the base of seed plants through a premature stop codon. *DIV* homologs often have a generic expression across *G. biloba*, but the expression of multiple *RAD* orthologs is significantly higher in ovules. This suggests a possible conserved role of *RAD* genes in seeds/ovules in angiosperms and gymnosperms.

## Results

### *DIV* genes have an elaborate history of duplication and loss

Bayesian phylogeny of *DIV* and *RAD* genes revealed that vascular plants have three clades of *DIV* genes: *DIV-A* (p = 0.97), *DIV-B* (p = 0.68), and *DIV-C* (p = 0.8) (Fig. 2). This suggests that the *DIV* gene in the common ancestor of vascular plants underwent two rounds of duplication, the first round generating *DIV*-*A* and the common ancestor of *DIV-B* + *DIV-C* (p = 0.94). The second round of duplication generated *DIV-B* and *DIV-C* (p = 0.94). Further, there have been duplications specific to the seed plants in the clades *DIV*-*A* and *DIV-B* (Fig. 2).

**Fig. 2.**
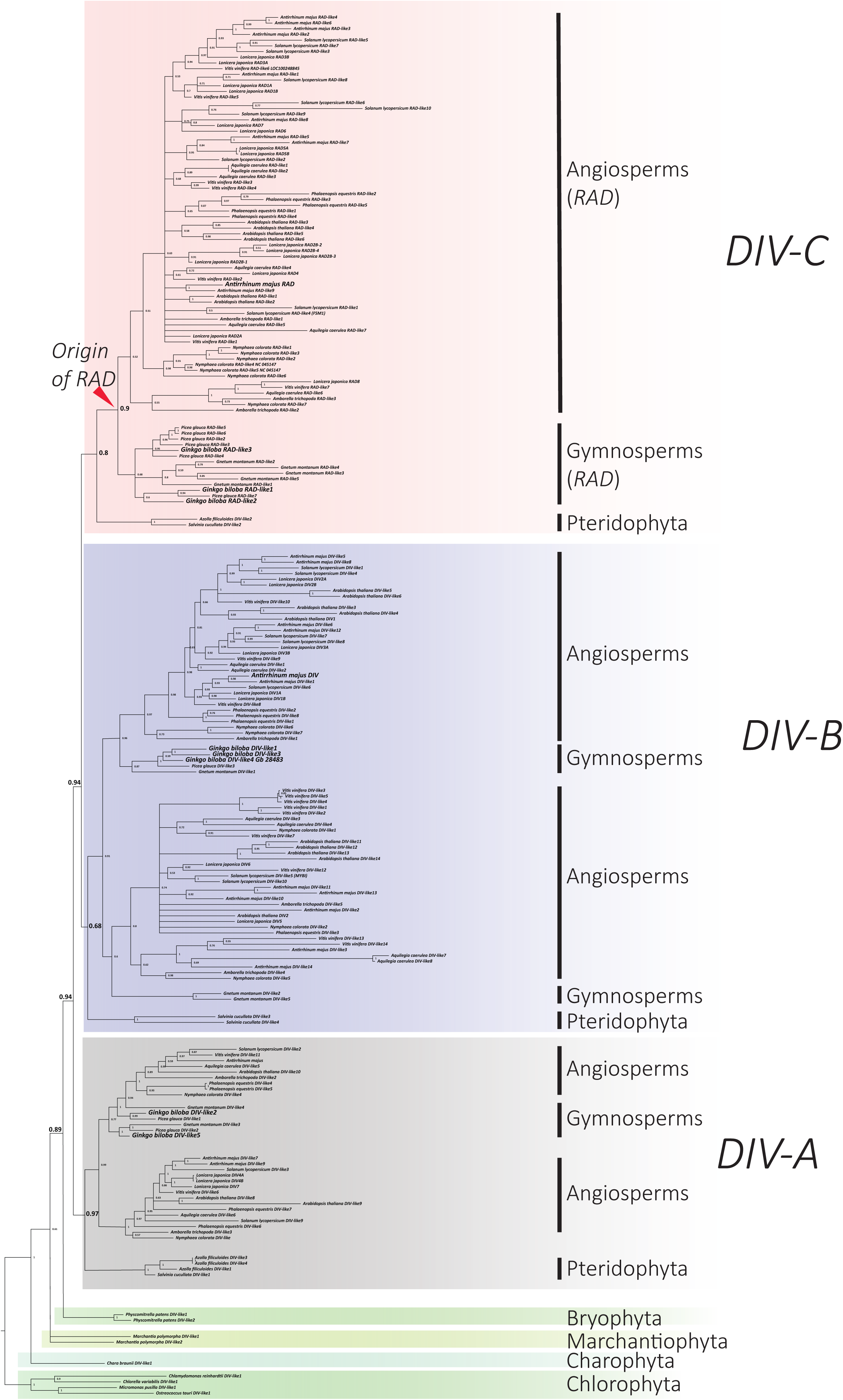
Bayesian phylogenetic tree of *DIV* and *RAD* genes in green plants based on nucleotide sequences. Posterior probability presented at nodes.

### Likely single origin of RAD-DIV interaction

The Bayesian phylogenetic tree of *DIV* and *RAD* genes (Fig. 2) provides strong evidence that *DIV* genes are ancestral to *RAD* genes because *RAD* genes are nested within *DIV* genes.

Further, these data support a single origin of *DIV* genes and a single origin of *RAD* genes, as opposed to multiple origins through parallel gain/loss of MYB2 domains (scenario A, Fig. 1). Though RAD evolved once, it is not impossible that RAD-DIV interactions evolved multiple times; testing such a hypothesis would require technology for wide-scale genetic manipulation and protein-protein interactions in both gymnosperms and angiosperms. Such experiments are currently not feasible. A parsimonious corollary of scenario A in Fig. 1 would be that the RAD-DIV competitive interaction likely originated once, at the base of gymnosperms + angiosperms. An additional line of evidence in favor of a single origin of RAD-DIV interaction is that in both known cases of RAD-DIV interactions (Machemer *et al*., 2011; Raimundo *et al*., 2013), the DIV component is from DIV-B clade (Fig. 2). In any seed plant genome, the *DIV-B* clade would be the closest relative of *RAD* genes (which are modified *DIV-C* genes). This would suggest that when RAD first evolved at the base of seed plants, it competed with its closest relatives, the DIV-B clade proteins, because these RAD and DIV-B proteins had similar protein sequences and expression patterns.

### *RAD* genes are modified *DIV-C* genes

Bayesian phylogenetic inference revealed that *RAD* genes are nested within *DIV* genes and are a part of the *DIV-C* clade with strong support (p = 0.9) (Fig. 2). *RAD* from gymnosperms and angiosperms are sisters to each other. Our phylogenetic analyses indicate that *RAD* evolved from a *DIV-C* like ancestor at the base of seed plants. *RAD* genes are present only in seed plants, and are a clade sister to pteridophyte *DIV-C* genes. To dissect the molecular processes that were associated with the modification of *DIV-C* into a *RAD* gene, we aligned *DIV* genes from ferns with all the *RAD* genes from broadly representative gymnosperms (*Ginkgo biloba*, *Picea glauca*, *Gnetum montanum*) and angiosperms (*Amborella trichopoda*, *Aquilegia caerulea*, *Antirrhinum majus*) (Fig. 3A). We found that *RAD* genes are truncated compared to *DIV-C* genes (Fig. 3B).

**Fig. 3.**
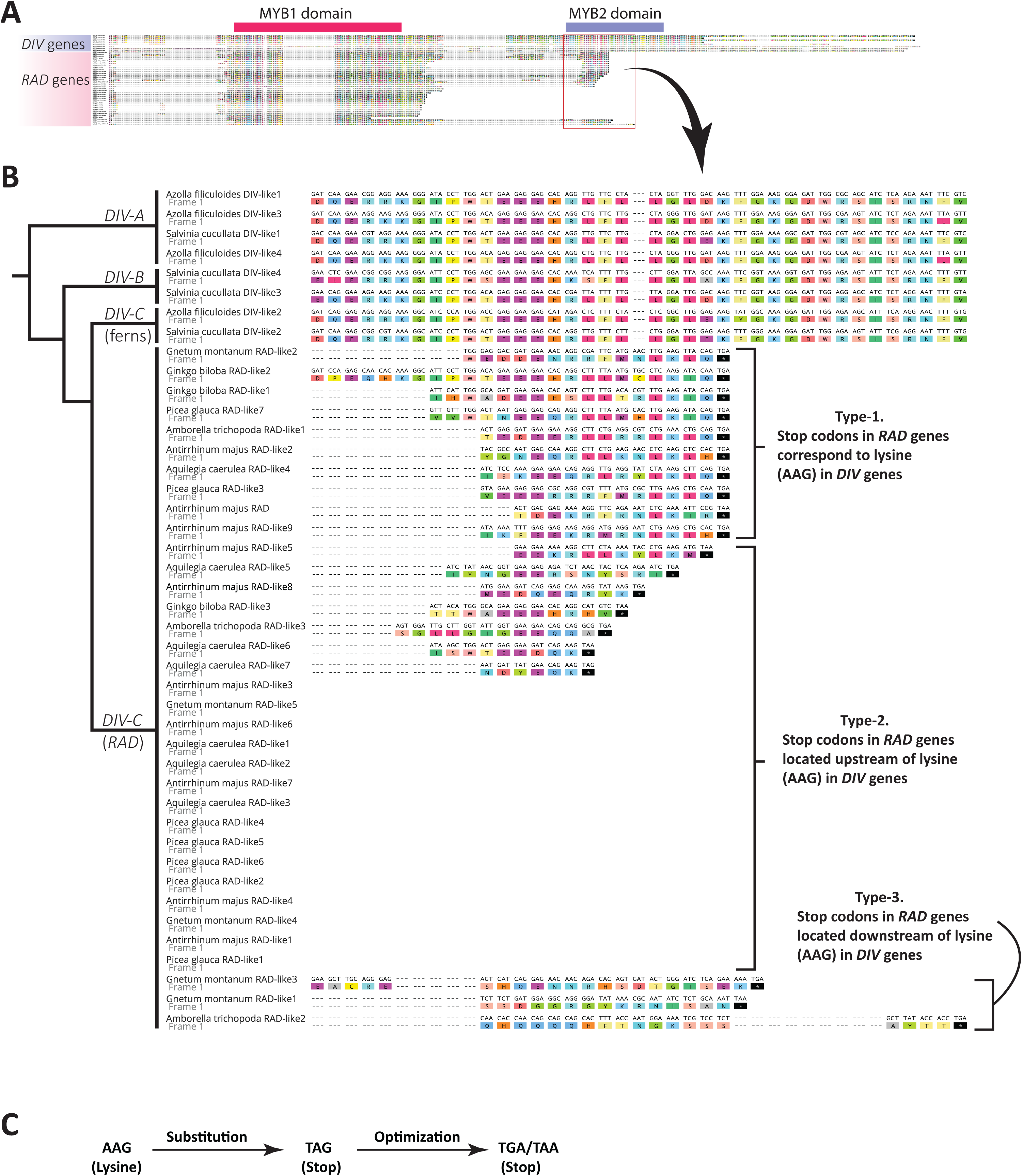
*RAD* genes evolved from *DIV* genes through an early stop codon. **A.** Alignment of *DIV* genes from ferns with all the *RAD* genes from representative gymnosperms (*Ginkgo biloba*, *Picea glauca*, *Gnetum montanum*) and angiosperms (*Amborella trichopoda*, *Aquilegia caerulea*, *Antirrhinum majus*). Nucleotide sequence along with translation displayed. Relative positions of MYB1 and MYB2 domains are designated. **B.** A portion of the alignment from A (boxed region) showing the beginning of the MYB2 domain. A simple phylogenetic tree displays the relationship among *DIV-A*, *DIV-B*, and *DIV-C* (including *RAD*) clades. Based on the location of stop codons, RAD genes can be of three types. Type-1, where the stop codons in *RAD* genes correspond to lysine (AAG) in *DIV* genes (this is likely the ancestral state for *RAD* genes). Type-2, where stop codons in *RAD* genes are located upstream of lysine (AAG) in *DIV* genes. Type-3, where the stop codons in *RAD* genes are located downstream of lysine (AAG) in *DIV* genes. **C.** *RAD* genes evolved from *DIV*-*C* genes through a premature stop codon. We postulate that the AAG codon in fern *DIV-C* clade genes (sister to *RAD*) became the stop codon TAG through substitution in the first position, and then to other stop codons (TGA, TAA) by further changes. The stop codon TAG is known to evolve into other stop codons for various evolutionary reasons.

*RAD* genes have a premature stop codon relative to *DIV* genes, and this stop codon is located early in what would have been a *MYB2* domain. Earlier reports had suggested that *RAD* genes only have a *MYB1* domain (Corley *et al*., 2005; Barg *et al*., 2005; Machemer *et al*., 2011), but we find that some *RAD* genes also have a truncated, vestigial *MYB2* domain represented by the early amino acids of the *MYB2* domain. (Fig. 3B).

*RAD* genes are variable both in their stop codon usage and their length. (Fig. 3B). Based on the location of stop codons, *RAD* genes can be of three types. Type-1, where the stop codons of *RAD* genes correspond to lysine (AAG) in *DIV* genes (Fig. 3B). Type-2, where stop codons of *RAD* genes are located upstream of this lysine (AAG) in *DIV* genes (Fig. 3B). Type-3, where the stop codons in *RAD* genes are located downstream of this lysine (AAG) in *DIV* genes (Fig. 3B). We think that Type-1 genes represent an ancestral state relative to Type-2 because the latter are shorter, that is, were derived by further shortening of the coding sequence by removing most of the truncated, vestigial MYB2 domain. Type-3, on the other hand, likely represents autapomorphies. This is based on the evidence that Type-3 stop codons appear only in individual homologs or clades, and that the additional coding sequences do not align well with other *DIV* or *RAD* genes. This suggests that Type-1 *RAD* genes preserve the ancestral feature with the stop codon of the ancestral *RAD* gene corresponding to lysine (AAG) in *DIV* genes, and Type-2 and Type-3 represent derived states.

Based on the evidence that the stop codon (TGA) in Type-1 *RAD* genes corresponds to a lysine residue (K) coded by AAG in fern *DIV* genes (Fig. 3B), we postulate the following two-step evolution of the stop codons in *RAD* from a fern-like state. First, the AAG codon became the stop codon TAG through substitution in the first nucleotide position, and then was further mutated to other stop codons (TGA, TAA) (Fig. 3C). The stop codon TAG is known to evolve into other stop codons (Belinky *et al*., 2018). This is not surprising, given that the stop codon TAG is less abundant in plants because it is leaky, susceptible to being misread, and is otherwise non-optimal (Angenon *et al*., 1990).

### Expression of *G. biloba RAD* and *DIV* homologs

We surveyed the expression of *RAD* and *DIV* homologs in *G. biloba* with qRT-PCR with a focus on reproductive organs (See Fig. 4). Functions of *RAD* and *DIV* homologs have been studied largely in the context of petals—an organ that gymnosperms, including *G. biloba*, do not have. The *DIV-A* clade is sister to the rest of *DIV* genes, and is represented by two genes in the *G. biloba* genome: *GbDIVlike5* and *GbDIVlike2* (Fig. 2). The only significant difference in expression level of *GbDIV-like5* is between leaves and a certain stage of the male strobilus (higher in leaves, Fig. 5). The remainder of the pairwise comparisons for *GbDIV-like5* are not significantly different. For the other *DIV-A* homolog, *GbDIV-like2*, the Tukey’s post-hoc test did not detect significant differences among tissue types though the p-value from ANOVA was significant suggesting that there are no pairwise differences (Fig. 5). This suggests that other comparisons—generated by permuting or pooling categories but not necessarily relevant to this study—could be significantly different. We compared several such combinations with Scheffé’s test (Stat trek website, Berman) but found no significant difference (CV_i_>L_i_). We also performed pairwise comparison employing Fisher’s test which detected some pairwise differences (Sup. Fig. 1). Fisher’s test does not provide protection against inflated Type 1 error rates arising due to multiple comparisons. Therefore, we interpret the data conservatively as showing no pairwise difference.

**Fig. 4.**
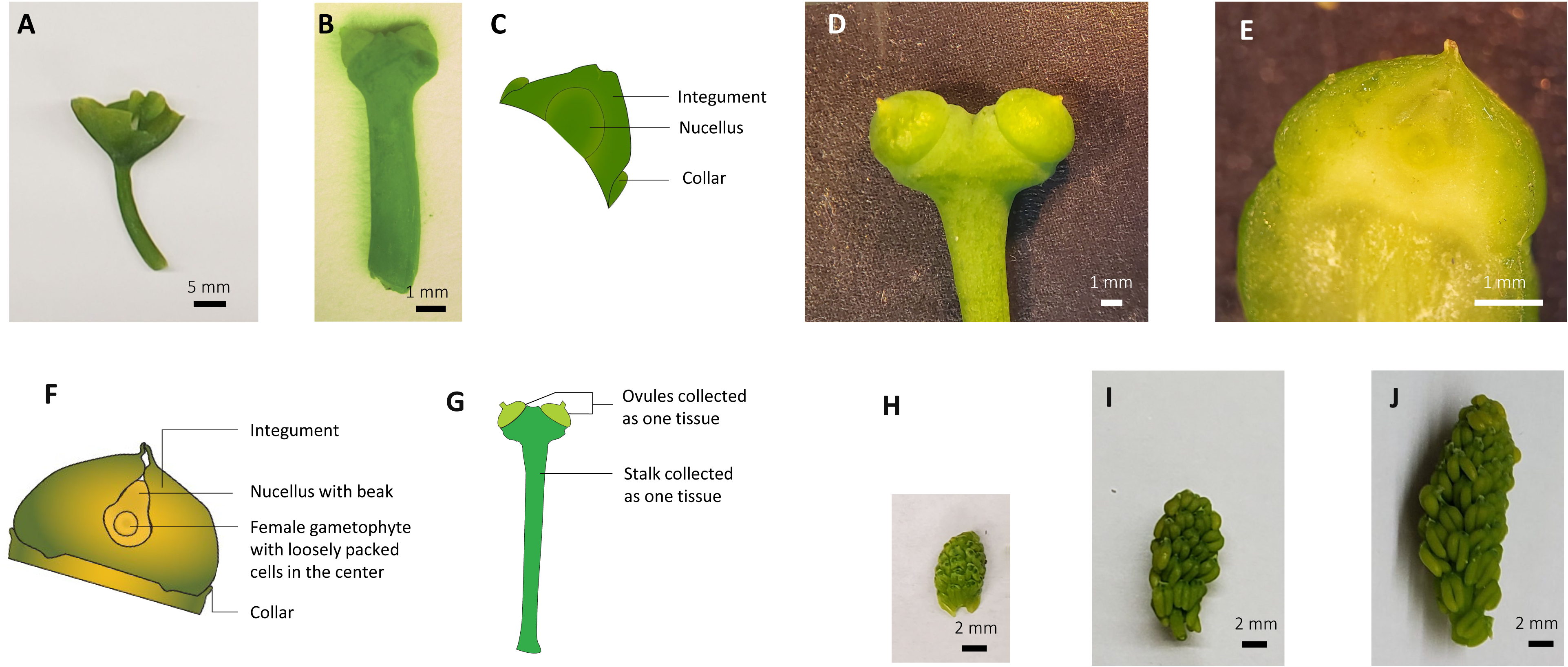
*Ginkgo biloba* tissues collected for qRT-PCR. **A.** A leaf from a female tree when the ovules are at stage-6 (*sensu* D’Apice *et al*., 2021) **B.** Female strobilus when the ovules are at stage-6 (*sensu* D’Apice *et al*., 2021). The height of the ovules (from the ovule-collar boundary to the tip of the integument) was *ca.* 0.6 mm. The diameter of the ovule was *ca.* 1 mm. The entire strobilus (stalk, ovules, and collars) was collected as a single tissue type. Tissue was collected on 12–13 April. **C.** Internal structure of ovule at stage-6 (*sensu* D’Apice *et al*., 2021). The female gametophyte is yet to develop. This unique anatomy is reflected in its gene expression pattern (Zumajo-Cardona *et al*., 2021), suggesting that it is a key developmental stage. **D.** Female strobilus when the ovules are at stage-9 (*sensu* D’Apice *et al*., 2021). The height of the ovules (from the ovule-collar boundary to the tip of the integument) was *ca.* 3 mm. The diameter of the ovule was *ca.* 4 mm. A part of the stalk is also visible. The ovules were collected as one tissue type, and the stalk was collected as another tissue type. Collected on 10–13 May. **E.** Longitudinal section of an ovule at stage-9 (*sensu* D’Apice *et al*., 2021) showing internal organization. **F.** A diagrammatic representation of the internal structure of the ovules at stage-9 (*sensu* D’Apice *et al*., 2021). The integument is starting to differentiate into three layers, the female gametophyte is undergoing cellularization, and pollination has likely happened. This unique anatomy is reflected in its gene expression pattern (Zumajo-Cardona *et al*., 2021), suggesting that it is a key developmental stage. **G.** Diagrammatic representation of how the tissue was collected from female strobilus when the ovules were at stage-9. The ovules were collected separately from the stalk. **H.** Male strobilus at length 0.7–0.8 cm, measured along the strobilus axis. Collected on 5 April. **I.** Male strobilus at length 1.3–1.4 cm, measured along the strobilus axis. Collected on 14– 15 April. **J.** Male strobilus at length 1.8–2.0 cm, measured along the strobilus axis. Collected on 14–15 April.

**Fig. 5.**
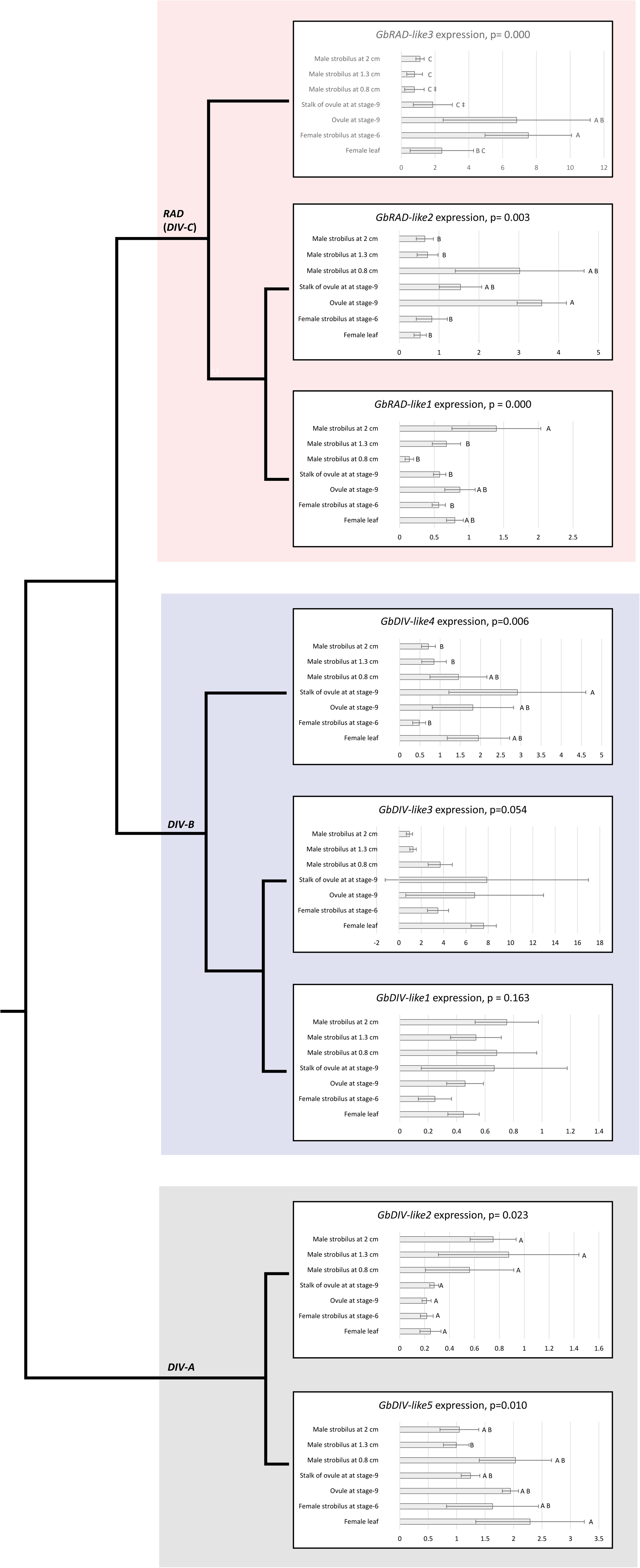
Relative expression levels of *RAD* and *DIV* genes in *Ginkgo biloba* determined by quantitative PCR. Error bars are standard deviations of samples. The p-values are from ANOVA (Fisher’s or Welch’s); letters at the tips are from post-hoc comparisons (Tukey’s or Games-Howell’s). Groupings from post-hoc comparisons are depicted by letters at the tip of the error bars; means that do not share a letter are significantly different. Expression of *GbRAD-like3* in tissues with the double dagger symbol (‡) had biological replicates whose one technical replicate displayed high Ct value and the other technical replicate did not amplify, suggesting a low expression level. In stalk of ovule, this was the case for all biological replicates; in male strobilus at 1.3 cm, this was the case for two of the five biological replicates. For *GbDIV-like2*, post-hoc test did not detect significant differences among tissue types despite the p-value from ANOVA being significant suggesting that there are there no pairwise differences (but other linear contrasts that are not relevant to this study).

The *DIV-B* clade includes orthologs of *AmDIV* which is involved in floral symmetry in *A. majus*. The *DIV-B* clade has two subclades in seed plants (Fig. 2). Each subclade has a gene whose protein is known to participate in a RAD-DIV interaction, though in different species: *AmDIV* (whose protein competes with AmRAD) (Raimundo *et al*., 2013), and *S. lycopersicum DIV-like5* (*SlDIV-like5*, i.e., *MYBI*, whose protein competes with SlRAD-like4, i.e., FSM1) (Machemer *et al*., 2011). *Ginkgo biloba* has three *DIV-B* clade genes: *GbDIV-like1*, *GbDIV-like3*, and *GbDIV-like4*—all three genes are from the subclade with *AmDIV* (Fig. 2). *Ginkgo biloba* has no representatives in the clade with *SlDIV-like5*, though another gymnosperm *Gnetum montanum* does (Fig. 2). In the expression data of two of the three *DIV-B* clade genes in *G. biloba*, *GbDIV-like1* and *GbDIV-like3*, ANOVA did not detect any significant differences (Fig. 5). For the third

*DIV-B* gene, *GbDIV-like4,* the stalk of the ovule at stage-9 has higher expression than whole strobili: whether male at later stages or female strobili at an earlier stage (Fig. 5).

*DIV-C* clade genes in *G. biloba* are represented by three *RAD* genes: *GbRAD-like1*, *GbRAD-like2*, and *GbRAD-like3* (Fig. 2). In *GbRAD-like1* all tissues have comparable expression levels, except that the male strobili at late stage (2 cm) have higher expression than most other tissues (Fig. 5). On the other hand, the remaining *RAD* genes in *G. biloba* have significantly higher expression in female tissues, especially in the ovules (Fig. 5). Particularly, the expression of *GbRAD-like3* is significantly higher in the ovules relative to the stalks that bear the ovule (Fig. 5). The expression of *GbRAD-like3* is also significantly lower in all stages of male strobili (Fig. 5).

## Discussion

### Hypothesized ancestral interaction involving RAD and DIV

DIV proteins are transcription factors with a diverse role in plant development: from flower symmetry (Galego & Almeida, 2002) to sugar homeostasis (Chen *et al*., 2019). Our phylogenetic analysis suggests that RAD proteins evolved from DIV via loss of a MYB domain. DIV proteins have two MYB domains: the protein-binding MYB1 domain and the DNA-binding MYB2 domain. RAD proteins have lost the MYB2 domain, and therefore act as repressors of DIV function by competing for the common protein targets of the MYB1 domain (competition described in (Machemer *et al*., 2011; Raimundo *et al*., 2013)). Two other kinds of competitive inhibition of DIV proteins by other MYB proteins are known, and in these cases the competitor competes for common *cis*-regulatory elements. (Chen *et al*., 2017, 2019). The first type include DIV proteins that can repress the transcription of genes normally upregulated by other DIV genes (Chen *et al*., 2017) and the second are LFG proteins (Chen *et al*., 2019). We have previously demonstrated that LFG proteins are truncated homologs of DIV proteins missing the MYB1 domain and that they evolved at the base of green plants (preprint: Sengupta & Howarth, 2025). Based on known interactions we have proposed a larger regulatory network that involves DIV, DRIF, LFG, and 14-3-3 proteins (preprint: Sengupta & Howarth, 2025).

It is possible that the network involving AmRAD and AmDIV (or their homologs) is much larger and includes the following: upregulator DIV, repressor DIV, RAD, DRIF, LFG, and 14-3-3 proteins (Fig. 6). In such a network, the function of AmDIV (or a homolog with upregulatory function) would be negatively regulated by RAD that competes for DRIF. An unknown repressor DIV protein (with both MYB1 and MYB2 domains) may compete with AmDIV for the same *cis*-regulatory site by employing its MYB2 domain (Fig. 6). This is based on the evidence from two DIV-A clade proteins, AtDIV-like10 and AtDIV-like8, that have both MYB1 and MYB2 domains, that compete for the same *cis*-regulatory sites (Chen *et al*., 2017). A similar competitive interaction may exist between AmDIV with another DIV homolog (or for various other pairs of DIV proteins). The protein-binding MYB1 domain of this unknown repressor DIV may bind to another protein, possibly a DRIF protein, because MYB1 domains are known to bind to DRIF proteins (Machemer *et al*., 2011; Raimundo *et al*., 2013). An unknown LFG protein (that only has the MYB2 domain, but no MYB1 domain) may also compete with AmDIV for the same *cis*-regulatory sites (Fig. 6) similar to the competition between the DIV protein OsMYBS1 and the LFG protein OsMYB2 (Chen *et al*., 2019). The amino acid motif in OsMYBS2 that binds to 14-3-3, including the phosphorylation site (serine), is widely conserved among LFG proteins (preprint: Sengupta & Howarth, 2025). This LFG protein may also bind to a 14-3-3 protein using a phosphorylated serine residue (Fig. 6).Interaction between 14-3-3 proteins and LFG proteins has been demonstrated in rice (Chen *et al*., 2019), and in soybeans (Dhaubhadel & Li, 2010; Li & Dhaubhadel, 2012; Dong *et al*., 2016).

**Fig. 6.**
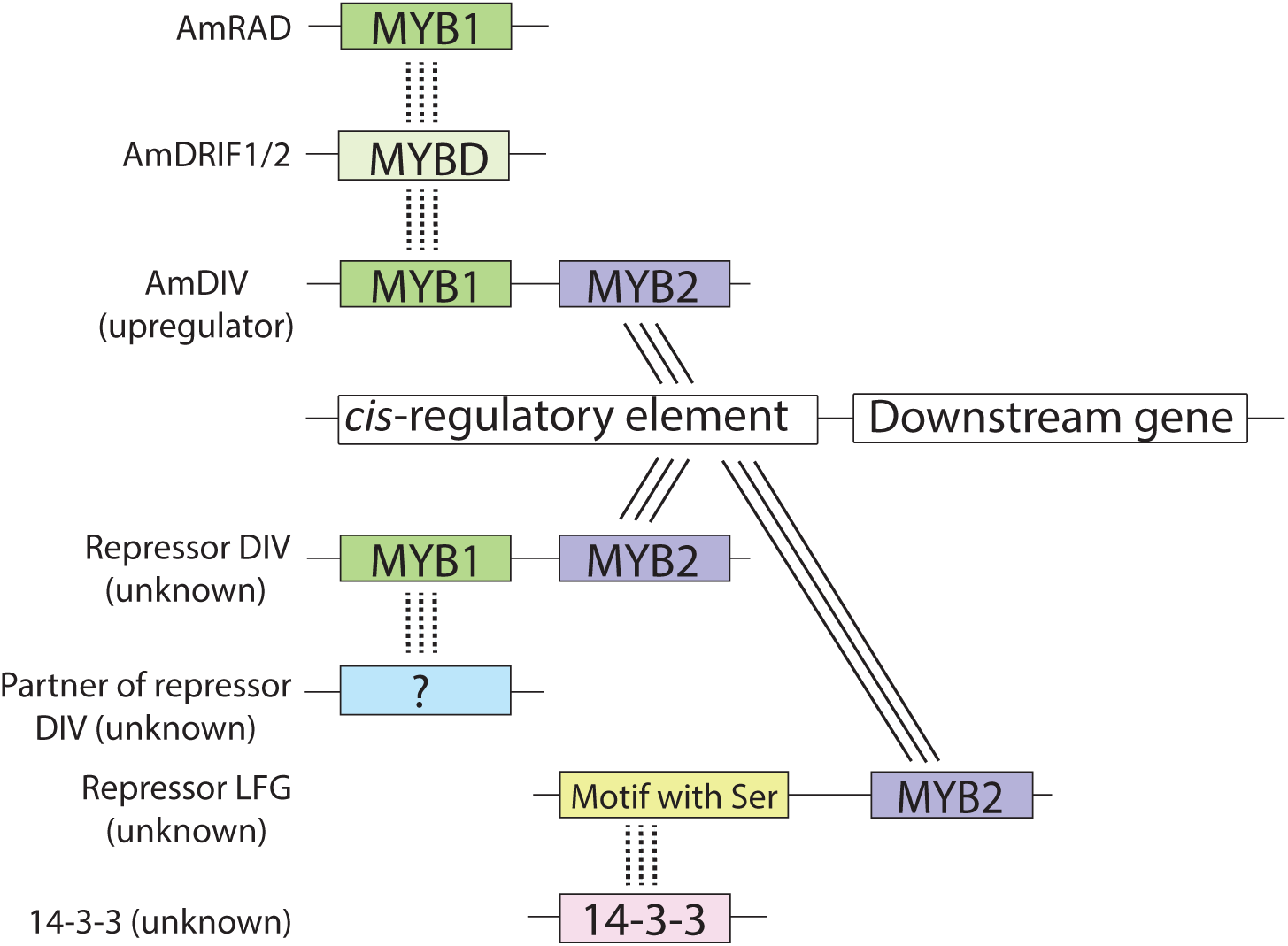
A model representing regulatory interactions of AmDIV that have been previously reported or are predicted in this work. We hypothesize that similar interactions exist for homologs of AmDIV. The transcription factor AmDIV can upregulate a target gene by binding to this gene’s *cis*-regulatory site (based on Raimundo *et al*., 2013). An unknown repressor DIV protein (with both MYB1 and MYB2 domain) may compete with AmDIV for the same *cis*-regulatory site (based on Chen *et al*., 2017) using the MYB2 domain, this repressing transcription. The protein-binding MYB1 domain of this unknown repressor of DIV may bind to another protein, possibly a DRIF protein (based on Machemer *et al*., 2011; Raimundo *et al*., 2013), but such partner of repressor DIV is currently unknown. An unknown repressor LFG protein (that only have MYB2 domain, but no MYB1 domain) may also compete with AmDIV (based on Chen *et al*., 2019) for the same *cis*-regulatory site. This LFG protein may bind to an unknown 14-3-3 protein using a phosphorylated serine (ser) residue (based on Chen *et al*., 2017). AmRAD represses AmDIV activity by competing for AmDRIF (Machemer *et al*., 2011). Protein-DNA interaction depicted by solid lines, protein-protein interactions by dotted lines.

All these genes in the hypothesized multi-gene network described above, except *RADIALIS*, are widely distributed in green plants, suggesting that an interaction among the following is possibly ancestral to green plants: upregulator DIV, repressor DIV, DRIF, LFG, and 14-3-3 proteins. With the evolution of RAD in seed plants, the ancestral interaction would have been expanded to include the competitive post-translational regulation of DIV by RAD. Given that several components of this hypothesized interaction are already known in *A. majus*, this species will be ideal to test for the remainder of the interactions. The ancestral function of this multi-protein network is not known.

### Hypothesized ancestral function of RAD and DIV

RAD evolved at the base of seed plants: they are truncated regulators of DIV, at least in model flowering plants. The origin of such competitive inhibitors have been considered to be associated with rapid morphological changes and increased complexity (Floyd *et al*., 2014). A similar competitive regulatory network has been described from vascular plants that involves regulation of C3HDZ function by their truncated relatives ZPRs (Floyd *et al*., 2014). Our data suggest that RAD evolved at the base of seed plants after their divergence from pteridophytes. Several morphological novelties evolved in this line leading to extant gymnosperms and angiosperms (Doyle, 2017). These include vegetative features (e.g., secondary xylem, cork cambium), male reproductive features (e.g., microspores as opposed to only homospores), and several features associated with female organs (Doyle, 2017). A major evolutionary innovation of the seed plants is the eponymous structure, the seed, and the structures associated with it.

Testing gene function in ovules through transgenics is unrealistic in most gymnosperms, including *G. biloba*, that take decades to reach reproductive maturity. However, we tested the expression of *RAD* and *DIV* genes in *G. biloba* by employing qRT-PCR to identify the potential functions of these genes. *DIV-A* genes in *G. biloba* are expressed at comparable levels in most tissues (Fig. 5). The *DIV-A* genes in *A. thaliana* are widely expressed across the plant (Sup. fig. 2). The *Physcomitrella patens DIV* genes that diverge immediately before the divergence of *DIV* genes in vascular plants are also expressed widely across the plant (Fernandez-Pozo *et al*., 2020). This suggests that the ancestral *DIV* gene at the base of vascular plants had a generic role in development. Within *DIV-B* clade genes (*AmDIV* orthologous group) two of the three *G. biloba* genes from this clade are expressed broadly across developmental stages and tissue types (Fig. 5). However, one ortholog, *GbDIV-like4*, has higher expression in ovules and stalks of ovules at stage-9 relative to some other tissues.

The expression pattern of *DIV-A* clade genes in *G. biloba* is consistent with the expression of *DIV-A* clades genes in flowering plants (Sup. Fig. 2). The expression pattern of the two *G. biloba DIV-A* genes suggest (Fig. 5) that the expression of the ancestral *DIV-A* clade gene was likely across the plant, and that it functioned as a generic developmental regulator (as opposed to defining particular organs).

The expression pattern of *DIV-B* clade genes in *G. biloba* is consistent with the expression of *DIV-B* clades genes in flowering plants. Two of the three *DIV-B* clade genes in *G. biloba* (*GbDIV-like1* and *GbDIV-like3*) are expressed widely across the tested tissue without any significant differences (Fig. 5). *AmDIV*, which is a *DIV-B* clade gene that defines the ventral side of *A. majus* flowers, is not expressed only in the ventral side, but in all floral organs (Galego & Almeida, 2002), and also in vegetative tissue (Sengupta & Hileman, 2022). The dorso-ventral nature of *AmDIV* function is derived from the fact that its competitor *AmRAD* is expressed only in the dorsal side of the flower and the AmRAD protein competes with the AmDIV protein in the dorsal region (Corley *et al*., 2005; Raimundo *et al*., 2013).The third *DIV-B* gene, *GbDIV-like4,* has higher expression in the stalk of the ovule at stage-9 than whole strobili: whether male at later stages or female strobili at an earlier stage (Fig. 5. This expression pattern is similar to that of *AmDIV* and *SlDIV-like5* that are expressed broadly across the plant but may have higher expression in some stages of female organ development (Sengupta & Hileman, 2022). The expression pattern of *DIV-B* clade genes in *G. biloba* is consistent with those from *A. thaliana*: most are expressed widely across the plant, but some (*AT3G10590*, *AT5G01200*) are also upregulated in fruits (Sup. fig. 2).

The *DIV-C* clade in seed plants is represented by *RAD* genes. The expression of two of the three *AmRAD* orthologs in *G. biloba* is higher in female tissues. Particularly, the expression of *GbRAD-like3* is significantly higher in the ovules relative to the stalks that bear the ovules, and is also lower in male strobili at all developmental stages. This expression pattern suggests that the ancestral function of *RAD* was likely a feminizing factor and it was involved in ovule development. *RAD* genes have been demonstrated to have a feminizing role (leading to the development of hermaphrodite flowers instead of male flowers) in the flowering plant *Diospyros* (Masuda *et al*., 2022) and this function has been hypothesized to be an ancestral function (as opposed to neofunctionalization) (Bergero, 2022). Expression data from divergent angiosperms also suggest that *RAD* genes are upregulated in flowers with female organs: *Schisandra chinensis* from Schisandraceae (Cheng *et al*., 2023) and *Plukenetia volubilis* from Euphorbiaceae (Fu *et al*., 2018). This is consistent with the expression of the *RAD* gene *At2g21650* in *A. thaliana* that is upregulated in carpels (Sup. fig. 2).The unit of female organ in *G. biloba* and angiosperms are structurally different, and their homology/homoplasy is currently ambiguous. What structure in *G. biloba* should be considered as a unit of female organ? There is debate about whether the female organ in *G. biloba* is the ovule, or the entire female reproductive shoot (which consists of two ovules and a stalk—including the collars around the ovules). Except for the ovules, which are clearly homologous between angiosperms and gymnosperms, the homology for the rest of the tissue is not clear between *G. biloba* female reproductive shoots and angiosperms carpels (Douglas *et al*., 2007; Gonçalves, 2021). The angiosperm carpel can be interpreted as a modified megasporophyll or as a modified rachis (discussed in Doyle, 2008). It is possible that the ancestral *RAD* gene had two functions: ovule development and overall female identity. The function in ovule development is likely conserved between angiosperms and *G. biloba.* If an ovule is interpreted as the female organ in *G. biloba* (instead of the whole female reproductive shoot), then the second, feminizing function of *RAD* was possibly associated with a transfer of identity in angiosperms where the identity of the female organ shifted from ovules (in gymnosperms) to carpels (in angiosperms). This would happen in the following two steps. First, ectopic expression of *RAD* would occur in the gymnosperm tissue homologous to carpels (either the rachis or the leaves, depending on interpretation), then these structures would evolve into modern day carpels.

The role of *RAD* in carpel development has not been tested broadly, but there is some evidence that *RAD* genes likely have such a role, at least in the pericarp (of *Solanum lycopersicum* or tomatoes, Machemer *et al*., 2011) and in the placenta (of *A. majus*, Li Feifei *et al*., 2023). However, these studies are based on over-expression of *RAD* genes, and are difficult to interpret because in over-expression mutants the interesting phenotypes may appear in parts of the plant where the gene is not natively functional. This indeed was the case when *AmRAD* was over-expressed in *A. majus*: the dorsal side of the flower (where the endogenous gene is expressed) showed no change, but the ventral part of the flower (where the endogenous gene is not expressed), showed a change in phenotype (Cui *et al*., 2010). So, it is possible that the function of RAD homologs in *S. lycopersicum* and *A. majus* is not in the regions affected in the over-expression mutants (pericarp or placenta), but in the ovules. This would be consistent with the high expression of *G. biloba RAD* genes in ovules.

There is no direct evidence implicating *RAD* genes in ovule development, primarily because it has not been explicitly tested with functional studies. It is important to note that high expression of *RAD* homologs in ovules has been reported in Lamiales (Zhong & Kellogg, 2015). High expression of *RAD* in female organs has been reported from a variety of flowering plants (though not always parsed into ovules vs. non-ovule parts of the organ) including magnoliids (Madrigal *et al*., 2019), monocots (Jain *et al*., 2007; Valoroso *et al*., 2017), lamiids (Zhong & Kellogg, 2015; Sengupta & Hileman, 2022), and rosids (Baxter *et al*., 2007; Fasoli *et al*., 2012). Such widespread expression of *RAD* in female organs across flowering plants, in addition to the functional evidence from *S. lycopersicum* (Machemer *et al*., 2011) and *Diospyros* (Masuda *et al*., 2022), has been considered to be an indication that *RAD* has a widespread role in female organ development in angiosperms (Sengupta & Hileman, 2022; Bergero, 2022). Our expression data suggest that a similar function likely exists in gymnosperms, and by corollary, that the ancestral function of *RAD* was in the development of female organs of seed plants.

The expression pattern of *RAD* in angiosperms and *G. biloba* is similar to that of the *AGAMOUS* (*AG*) clade of MADS-box genes. In angiosperms, homologs of *AG* are expressed in and provide identity to both carpels (Bowman *et al*., 1989, 1991) and to ovules (Bowman *et al*., 1991; Pinyopich *et al*., 2003). The orthologs of *AG* from *G. biloba* (D’Apice *et al*., 2022), and another early diverging gymnosperm *Cycas edentata* (Zhang *et al*., 2004), are expressed in ovules, suggesting a conserved function of *AG* in ovule development across seed plants. This expression pattern—in carpels of angiosperms, and in ovules of both gymnosperms and angiosperms —is similar to that of *RAD*. It is possible that the genetic network involved in ovule development was co-opted during the evolution of carpels, and that both *AG* and *RAD* are involved in this network.

## Conclusion

RAD-DIV competitive interaction is critical to the development of bilaterally symmetric flowers in Lamiales. A similar interaction is also involved in fruit development in *S. lycopersicum* (Solanales). We provide evidence that *RAD* genes evolved once, suggesting that the RAD-DIV competition seen in distant species also evolved once, and was inherited/modified from the ancestral seed plant. Further, we provide preliminary evidence that the ancestral function of this interaction in seed plants was likely in female organ development, especially, of the ovules. Thus, a major event in the history of plants—the origin of the seed habit—was possibly associated with the origin of *RAD* genes.

## Materials and methods

### Identifying homologs

We identified the homologs of *DIV* and *RAD* genes by performing nucleotide BLAST and tBLASTX (Altschul *et al*., 1990) searches on published sources. When the coding sequence had not already been determined or predicted by previous authors, or when the previously reported coding sequence did not match the expectations from the closest homologs, we determined the coding sequence by alignment with other genes. The gene *Picea glauca RAD-like1* (clone GQ0207_B03 BT103128) was not included in the study because it has a very short sequence that interferes with downstream phylogenetic analysis. All homologs studied in this work and their sources are listed in Supplementary table 1. The unmodified, unaligned coding sequences are presented in Supplementary file 1.

### Phylogenetic analysis

The data was aligned using the amino acid coding sequence in Geneious Prime (Kearse *et al*., 2012) using MAFFT v7.450 (Katoh *et al*., 2002). We manually improved the alignment and removed all columns with 70% or more gaps.

We used Bayesian analyses to estimate the phylogenetic history of *DIV* and *RAD* using MrBayes 3.2.7a (Ronquist *et al*., 2012) available at CIPRES (Miller *et al*., 2010) [www.phylo.org]. We employed minimally informative (also called uninformative or flat) priors and ran the program for 10 million generations. The nexus alignment file with the Bayesian command block is presented in the Supplementary file 2.

### Tissue sampling

We collected vegetative and reproductive tissue from male and female *G. biloba* trees growing near St John’s University (Queens, NY), in Queens Botanical Garden (Queens, NY), and at Levittown Public Library (Levittown), NY. We collected male and female reproductive tissue at various stages of development and also collected leaves from female trees (Fig. 4). The developmental stages of the female structures were determined from published descriptions (D’Apice *et al*., 2021; Zumajo-Cardona *et al*., 2021), and with advice from Dr. P. L. Uniyal (University of Delhi) and Dr. S. Moschin (University of Padova).

### RNA extraction and Real-time PCR

Our initial tests with RNA extraction and qRT-PCR demonstrated that the extract had PCR inhibitors because decreasing the template amount in the PCR would elevate the estimated expression level. We tested various combinations of RNA extraction kits, cDNA synthesis kits, one- and two-step qPCR kits, and PCR additives. We tested the following RNA extraction kits: E.Z.N.A. Plant RNA Kit (R6827-01, from Omega Bio-tek, Norcross, Georgia, USA), RNeasy Plant Mini Kit (74903, from Germantown, Maryland, USA), MagMAX Plant RNA Isolation Kit (A33784, from Applied Biosystems and ThermoFisher Scientific, Waltham, MA), and Plant/Fungi Total RNA Purification Kit (25800, from Norgen Biotek Corp, Thorold, ON, Canada). The performance of RNA extraction kits varied among tissues. We tested one-step qRT-PCR kits (SsoAdvanced Universal SYBR Green Supermix, 1725270, from Bio-Rad, Hercules, California, USA; and qScript One-Step SYBR Green qRT-PCR Kit, 95087-050, from Quantabio, Beverly, MA, USA) and two-step qRT-PCR kits (Bullseye EvaGreen qPCR 2x Mastermix-ROX, BEQPCR-R, from Midwest Scientific, Fenton, Missouri, USA; and PerfeCTa SYBR Green FastMix, 84069, from Quantabio). We tested cDNA synthesis kits (qScript cDNA Supermix, 95048-100, from Quantbio; and PR1MA qMAX cDNA Synthesis Kit, PR2110, from Midwest Scientific). We also tested various additives/enhancers that are known to improve regular PCR (Grunenwald, 2008) with two-step qRT-PCR at the following final concentrations: dimethyl sulfoxide (DMSO, 5% v/v), glycerol (5% v/v), formamide (5% v/v), bovine serum albumin (BSA, 50 μg/μl), polyethene glycol-4000 (PEG-4000, 5% w/v), polyethene glycol-20,000 (PEG-20,000, 5% w/v, 901051-100ML-F, from Sigma Albuquerque, New Mexico, USA), silwet (0.5% v/v). We also tested lithium chloride based purification of RNA extract that is expected to remove PCR inhibitors from the extract (Zhang *et al*., 2021; ‘Technical Bulletin: #160. LiCl Precipitation for RNA Purification’).

Informed by these comparisons, we selected the following protocol of extraction and qRT-PCR. We extracted RNA from the male tissues and the stalk of the female strobilus using RNeasy Plant Mini Kit (with 100 μL of PEG-20,000 added to 1 ml of buffer RLT). For the remaining tissue types, we used E.Z.N.A. Plant RNA Kit with a similar addition of PEG-20,000 to the buffers NLT and RB. We did not clean the RNA extracts with lithium chloride because our tests showed no significant improvement in the quality of RNA. The best results for qRT-PCR were obtained when using two-step qRT-PCR without any PCR enhancers/additives.

For two-step qRT-PCR, we first generated complementary DNA (cDNA) from the RNA using qScript cDNA Supermix. We then performed qRT-PCR on the cDNA using Bullseye EvaGreen in CFX96 Touch Real-Time PCR Detection System (Bio-Rad). We used the gene *Ginkgo biloba HAS28* (*28 kDa heat-and acid-stable phosphoprotein*, Gb_13272) as the reference housekeeping gene because it has been demonstrated to be an appropriate reference gene in this species for comparison among various developmental stages and tissue types (Zhou *et al*., 2020). We determined the efficiency of the reactions and the primers using the DART-PCR algorithm that determines amplification efficiency from the slope of the amplification plot (Peirson *et al*., 2003). We measured relative gene expression using the Pfaffl method (Pfaffl, 2001, 2007). We confirmed that the primer pairs only amplified the correct target—we cloned PCR products of each primer pair into a sequencing vector and sequenced the amplicon. We performed statistical analyses using Microsoft Excel and Minitab (‘Minitab Statistical Software’)

In our qRT-PCR experiments, each tissue type was represented by three to five biological replicates; each biological replicate being from a different tree. Further, each biological replicate was represented by two technical replicates. To estimate whether the expression was significantly different among tissue types, we first performed Levene’s test of homogeneity of variance. For genes whose expression levels in different tissues had homogeneous variance, we performed Fisher’s classic one-way ANOVA, followed by Tukey’s post-hoc pairwise comparisons (when ANOVA had significant p-value). For the gene *Ginkgo biloba RADIALISlike2*, Levene’s test suggested non-homogeneity of variances. Therefore, we performed Welch’s one-way ANOVA for this gene, followed by Games-Howell’s post-hoc test.

## Acknowledgements

The authors thank Dr. P. L. Uniyal (University of Delhi) and Dr. S. Moschin (University of Padova) for help with identifying the stages of development of *G. biloba*. Devesh Shah (graduate student, St. John’s University) assisted with tissue collection. The authors thank Morgan Potter (Supervisor of Gardeners, Queens Botanical Garden) and the Levittown Public Library for help with collection.

## Author notes

ASG conceived of this project and developed it with advice from DGH. ASG performed all experiments and analyses. ASG prepared the first draft of the manuscript. ASG and DGH finalized the manuscript. All authors have read and approved the manuscript.

## Data availability

The data underlying this article are available in the article and in its online supplementary material.

## Ethics approval statement

All experiments were conducted in adherence to local, state, and federal regulations.

## Supplementary data

Supplementary file 1. The unmodified, unaligned coding sequences of genes used in this study. Supplementary file 2. Nexus alignment and MrBayes command block used for phylogeny.

Supplementary table 1. Source of *RAD* and *DIV* genes used in phylogenetic analysis.

Supplementary table 2. Primers for quantitative real-time PCR of *Ginkgo biloba DIV*, *RAD,* and *HAS28* genes.

## Supplementary figures

Supplementary fig. 1. Relative expression levels of *GbDIV-like2* compared with Fisher’s post-hoc test. Error bars are standard deviations of samples. The p-value is from Fisher’s ANOVA; letters at the tips are from Fisher’s post-hoc comparisons.

Supplementary fig. 2. Expression of *DIV* (and *RAD*) homologs in *Arabidopsis thaliana* and *Solanum lycopersicum.* The expression graphs are from The Bio-Analytic Resource for Plant Biology available at https://bar.utoronto.ca/ (Waese & Provart, 2017). Additional information for *At4g36570* was taken from Baxter *et al*. (2007).

