## Supplementary figures and images for "*RADIALIS* origin and expression suggest ancestral function in female organs of seed plants"

### Supplementary Figures.pdf

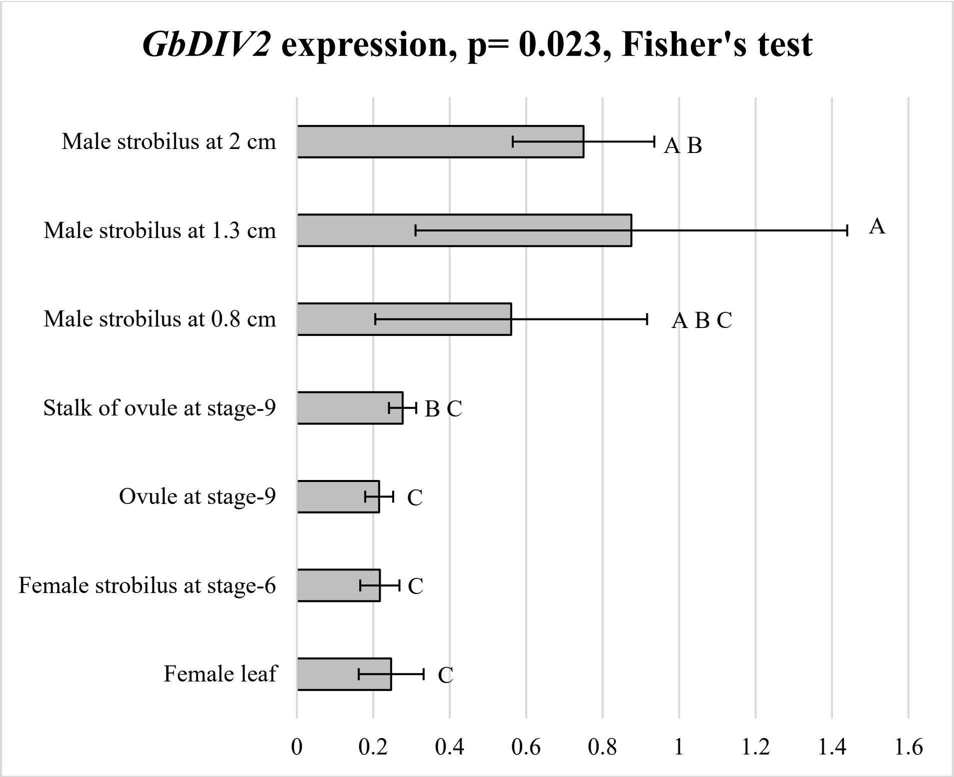

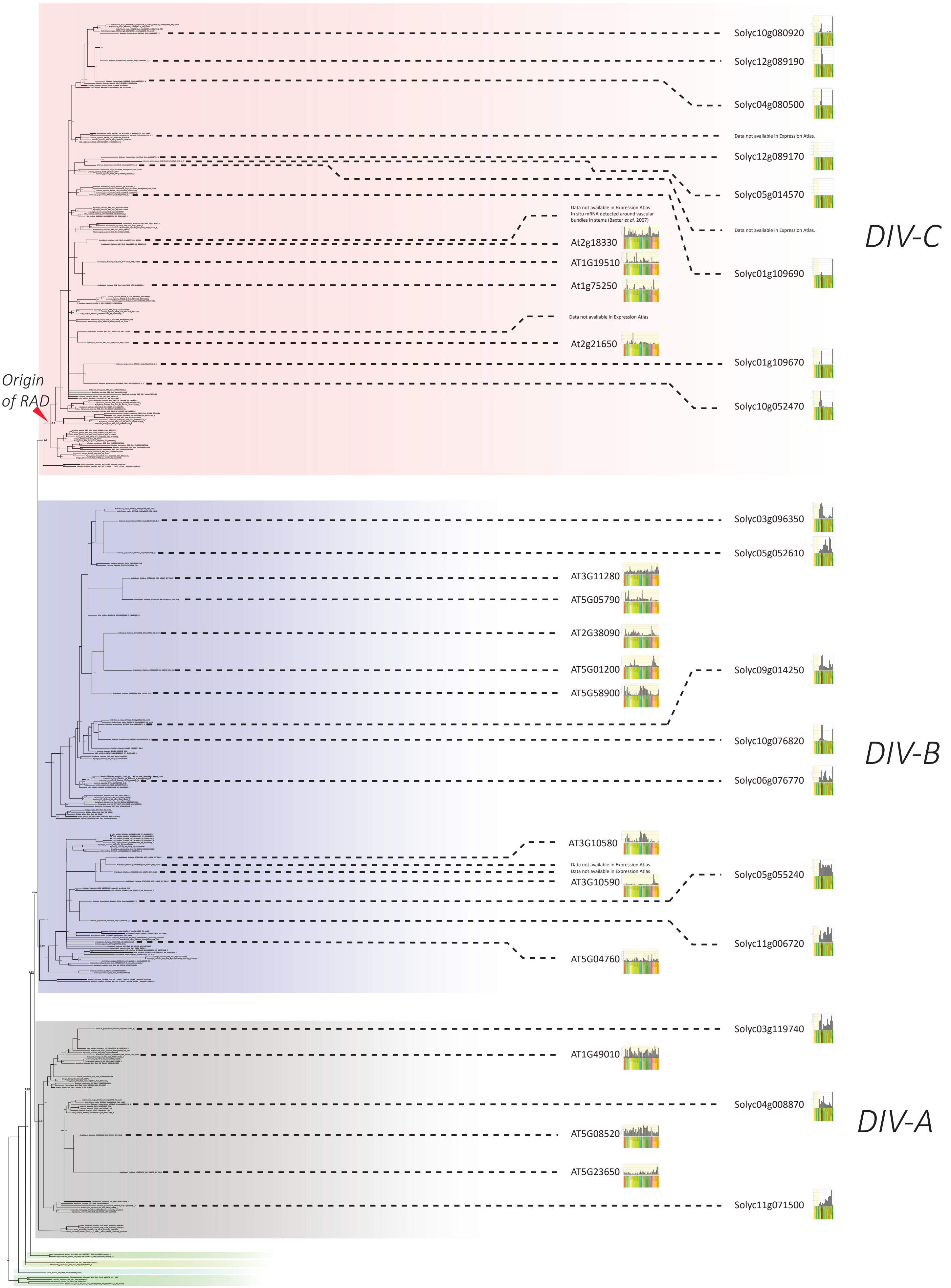
